# Engineering armoured *in vivo* CAR T cells through targeted delivery and transient mRNA gating

**DOI:** 10.64898/2026.03.10.710759

**Authors:** Kai Hu, Saket Srivastava, Biao Ma, Yajing Yang, Huimin Wang, Maurizio Mangolini, Jasvinder Hayre, Emily Souster, Rajesh Karattil, Aubin Ramon, Matthew Greenig, Pietro Sormanni, Shaun Cordoba, Shimobi Onuoha

## Abstract

*In vivo* generation of chimeric antigen receptor (CAR) T cells offers an off-the-shelf, scalable alternative to *ex-vivo* therapies, but is constrained by inefficiency, toxicity and durability. We report the first armoured in-vivo CAR T platform with tunable, transient activity, that enhances safety and efficacy.

Our platform utilises CD8 targeted lipid nanoparticles (LNP) to transiently deliver CAR and armouring membrane-bound IL12 mRNAs. Payloads utilize a T-cell-restricted (T-trex) strategy to constrain CAR and IL-12 expression to T cells, enabling potent antigen-dependent activity with spatio-temporal control.

Selective targeting of T-cells with a T-trex CAR showed enhanced expression. The addition of T–trex IL12 mRNA provides a tunable, localised amplification of CAR function, enhancing antigen-dependent cytotoxicity and cytokine production with robust tumour control.

These findings establish a novel dose-efficient armoured *in-vivo* CAR T approach and general framework for programmable immune cell engineering.

## Introduction

Chimeric antigen receptor (CAR) T cell therapies have demonstrated transformative efficacy in B cell malignancies and have more recently shown promise in selected immune-mediated diseases^1,2^. In oncology, durable tumour clearance has been achieved through deep and sustained elimination of target antigen–expressing cells, enabled by high effective CAR T cell doses and prolonged functional persistence following *ex vivo* manufacture^3^. However, despite their clinical impact, *ex vivo* CAR T cell therapies remain limited by complex and individualised manufacturing, high cost, restricted scalability, and treatment-associated toxicities, motivating the development of alternative approaches that enable broader access and greater programmability ^4,5^.

*In vivo* CAR T cell generation has emerged as a compelling strategy to overcome the limitations of *ex vivo* manufacturing by enabling off-the-shelf administration^6^. Multiple delivery modalities have been explored, including viral vector–based approaches and cell-targeted nanoparticle systems, each with distinct advantages and constraints^7,8^. Among these, targeted delivery of CAR-encoding messenger RNA (mRNA) using lipid nanoparticles (LNPs) has demonstrated particular promise due to its transient expression, improved safety profile and potential for scalable manufacturing^9^. Preclinical studies have shown that LNP-mediated CAR mRNA delivery can reprogram T cells *in vivo*, resulting in tumour control across cancer models and functional CAR activity in nonhuman primates^10^. More recently, early clinical feasibility has been reported using CD8-targeted LNPs delivering CD19 CAR mRNA, providing the first demonstration that *in vivo* CAR T cell generation can be achieved safely in humans^11^. Together, these studies establish important proof-of-principle for LNP-based *in vivo* CAR approaches across both malignant and non-malignant indications.

Despite this progress, current *in vivo* CAR strategies face fundamental constraints that limit their broader therapeutic impact. In contrast to *ex vivo* CAR T cell therapies, where sustained CAR expression can be achieved, CAR expression following *in vivo* mRNA delivery is inherently transient and current platforms require re-dosing every 2-3 days to sustain CAR expression ^10,11^. As a result, therapeutic efficacy is often limited by insufficient depth or duration of CAR activity, particularly in disease settings that require robust and sustained target cell clearance, such as high tumour burden malignancies or settings with antigen re-emergence^1,2^.Attempts to overcome these limitations through dose escalation are restricted by toxicity and the feasibility of highly frequent repeat administration to maintain CAR expression, a feature that is increasingly recognised as important for maintaining disease control and preventing relapse.

These challenges highlight a central problem in the design of *in vivo* CAR therapies: how to achieve durable biological efficacy without relying solely on escalating dose to prolong and maintain CAR expression. We hypothesised that overcoming this limitation would require a programmable system that enhances the functional potency of individual CAR-expressing T cells, rather than simply increasing the number of cells transfected or the amount of material delivered. In this context, strategies that lower the activation threshold of CAR T cells, restrict expression to defined immune subsets, and permit repeat dosing at low total exposure may be particularly well suited to both malignant and non-malignant disease settings.

Here, we describe a programmable *in vivo* CAR T cell platform that integrates three complementary engineering layers: (i) a CD8-targeted, spleen-tropic LNP delivery system, (ii) mRNA payloads that are selectively expressed on T cells, and (iii) transient, surface-tethered IL-12 armouring to locally enhance CAR T cell activation. By lowering the functional activation threshold of CAR-expressing T cells while maintaining tight control over cytokine exposure, this approach enables robust biological activity at doses compatible with repeat administration. Using a CD19 CAR as a model system, we demonstrate that this platform supports deep and sustained target cell depletion *in vivo* using ultra-low total CAR doses, establishing a generalizable framework for next-generation *in vivo* CAR T cell therapies applicable to both malignancies and immune-mediated diseases.

## Results

### Engineering of a novel armoured targeted LNP platform to facilitate dose-efficient in vivo CAR delivery

To enable dose-efficient and repeatable *in vivo* CAR T cell generation, we developed a delivery platform integrating targeted lipid nanoparticles, T-cell-restricted mRNA expression, and transient cytokine armouring (Fig. 1A). This architecture was designed to enhance the functional potency of CAR-expressing T cells while limiting systemic exposure, thereby reducing the total dose required to achieve robust biological activity.

**Figure 1.**
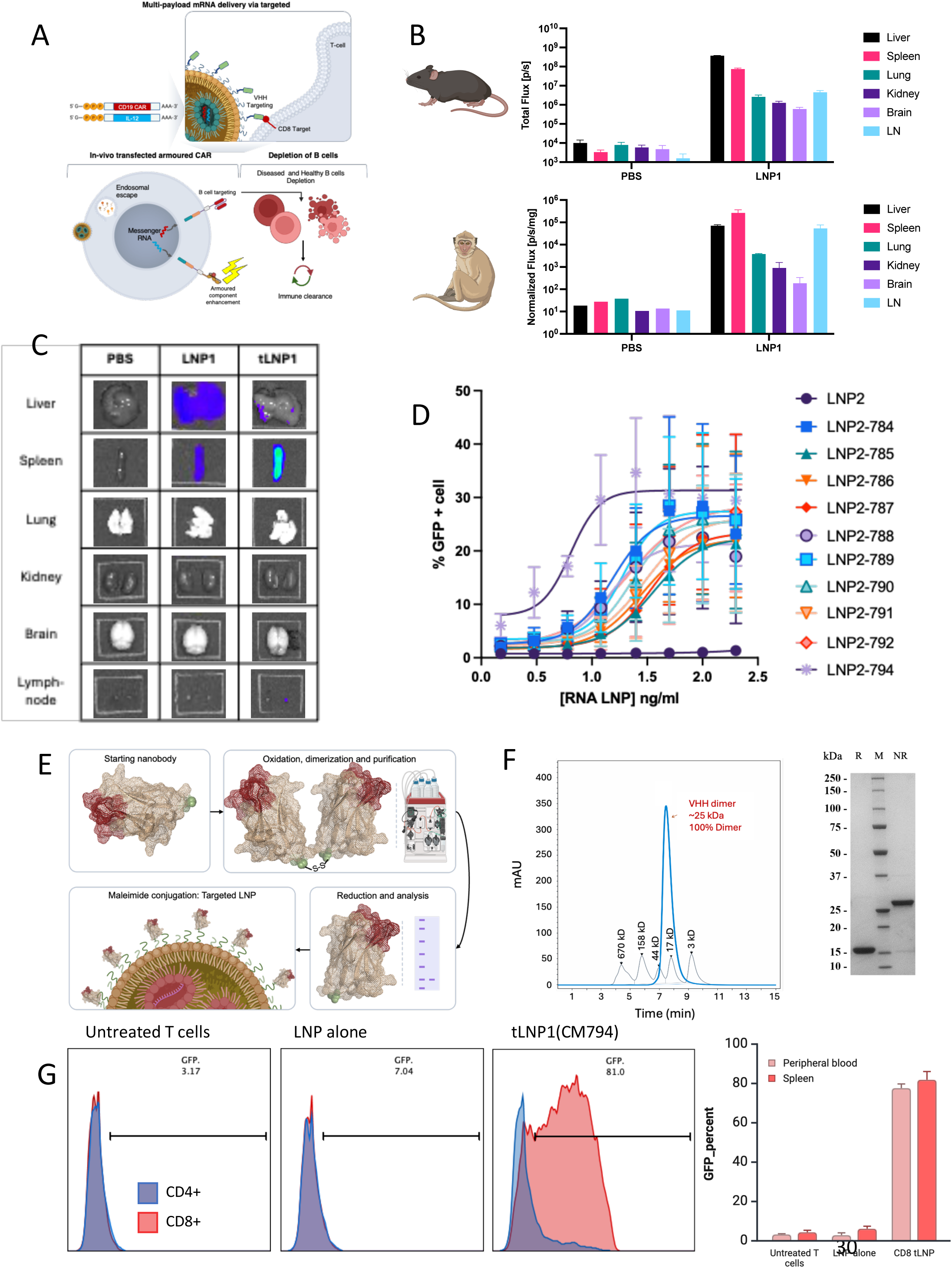
Targeted LNP platform enables specific and repeatable *in vivo* CAR delivery. A. Integrated platform schematic: Overview of the three-layer engineering strategy: multi-payload mRNA (encoding CAR and tethered IL-12) delivered via CD8-targeted LNPs to achieve selective T-cell reprogramming and B-cell depletion. B. Passive splenic tropism: Quantitative biodistribution analysis showing total flux (p/s) in mice (top) and normalized flux (p/s/mg) in cynomolgus macaques (bottom) following systemic administration of untargeted LNP1. Formulation LNP1 exhibits intrinsic spleen accumulation across species, though liver expression persists. C. Refinement of biodistribution via active targeting: Representative *ex vivo* bioluminescence imaging of major organs in mice treated with PBS, untargeted LNP1, or murine CD8-targeted LNPs (tLNP1). Targeted conjugation markedly increases spleen specificity and reduces hepatic signal^5^. D. Functional screening of humanized VHH candidates: Dose-response curves identifying CM794 (purple) as the lead binder for mRNA-driven protein expression in human T cells. Candidates were humanized using the AbNatiV dual-control strategy to balance humanness and nativeness. E. VHH manufacturing and conjugation strategy: Schematic illustrating the engineering of homodimeric VHH constructs (oxidation/dimerization) followed by controlled reduction to provide maleimide-accessible monomers for precise LNP surface decoration. F. Biophysical characterization of targeting ligands: Size exclusion chromatography (SEC) chromatogram confirming a single homodimeric peak (Approx 25 kDa, 100% dimer). Right: SDS-PAGE analysis of purified CM794 under reducing (R) and non-reducing (NR) conditions, confirming efficient dimerisation and reduction. G. Validation of CD8-specific targeting: Representative flow cytometry histograms from in-vitro PBMC manipulation(left) and quantitative analysis from in-vivo transfection (right) show selective association between CM794-tLNPs and CD8⁺ vs CD4⁺ T cells in human PBMCs and engrafted NSG mice. Specificity is maintained in both peripheral blood and splenic compartments.

We first evaluated whether LNP composition alone could support immune-cell-selective delivery *in vivo*. Screening of a panel of proprietary LNP formulations identified an LNP formulation (LNP1) with intrinsic spleen accumulation following systemic administration in both mice and cynomolgus macaques (Fig. 1B). However, despite identification of nanoparticles with favourable biodistribution, measurable expression in non-lymphoid tissue, primarily in the liver, was consistently observed, indicating that passive tropism alone was insufficient to achieve selective targeting of the desired immune effector cells.

To further enhance specificity, we explored active targeting strategies based on CD8 engagement. Conjugation of a murine CD8-targeting antibody to the LNP surface resulted in a marked increase in spleen specificity *in vivo* in mice compared with untargeted formulations (Fig. 1C), supporting the use of ligand-mediated targeting to refine biodistribution beyond that achievable through lipid composition alone.

To identify suitable CD8-targeting ligands for delivery to human T cells, a panel of humanised VHH-based antibodies were generated. To minimize the risk of anti-drug antibody (ADA) formation while preserving physicochemical stability, we employed a deep learning-based dual-control humanisation strategy^12^. This approach utilized the AbNatiV vector-quantized variational auto-encoder (VQ-VAE) to engineer humanising mutations into parental camelid-derived VHH frameworks^12^. By simultaneously scoring variants for *humanness* (likelihood of belonging to the human immune antibody repertoire) and *VHH-nativeness* (likelihood of belonging to stable camelid single-domain antibody repertoire), we prioritised candidates predicted to minimise immunogenicity without compromising structural integrity or biophysical suitability for LNP surface conjugation. A representative humanised design selected following functional screening is shown in Supplementary Fig. 1A. This dual-optimisation strategy avoids the trade-off between immunogenicity and stability that is often observed with conventional residue-frequency-based humanisation approaches. To generate more diversity, we ran both ‘Enhanced’ and ‘Exhaustive’ AbNatiV mutational sampling pipelines^12^, either limited to framework resurfacing (default behaviour), or allowing also mutations at buried positions or in the CDRs. This approach generated several humanised candidates, which exhibited a range of biophysical properties (Supplementary Fig. 1B), necessitating the development of a functional screening strategy capable of identifying ligands compatible with LNP conjugation and cellular uptake. A modular screening platform was therefore established, using covalent irreversible capture via molecular glue based universal LNPs, enabling rapid evaluation of multiple CD8-targeting binders in an LNP-associated context (Supplementary Fig. 1C).

Functional screening for internalisation and mRNA driven protein expression identified CM794 as a lead CD8-targeting ligand (Fig. 1D). Importantly, no binding to CD4⁺ T cells was observed across tested concentrations using cells from the same donors (Supplementary Fig. 1D), confirming CD8 specificity of the selected binder.

To enable reproducible and scalable conjugation of CM794 to LNPs, homodimeric VHH constructs were engineered to support controlled generation of conjugation-competent monomers (Fig. 1E). Biophysical characterisation confirmed that these constructs were fully homodimeric prior to reduction and could be efficiently and reproducibly reduced under mild conditions without detectable aggregation (Fig. 1F and Supplementary Fig. 1E). Analysis of antibody surface conjugation demonstrated controlled and reproducible decoration of the LNP surface across batches (Supplementary Fig. 1F), establishing reproducible ligand tethering as a manufacturable parameter.

Finally, CM794-conjugated LNPs were evaluated for functional targeting in human PBMCs *in vitro* and following engraftment into NSG mice. In PBMCs, targeted GFP LNPs demonstrated significantly greater expression selectivity in CD8^+^ T cells compared to other PBMC compartments (Supplementary Fig 1G). Upon systemic administration in mice, expression specificity in CD8^+^ T cells was observed in both splenic and peripheral blood compartments (Fig. 1G). Together, these data establish a targetable, manufacturable LNP platform capable of selectively delivering CAR mRNA to CD8⁺ T cells, forming the foundation for *in vivo* CAR T cell generation.

### RNA-level gating enables potent and durable T-cell-restricted CAR expression

Having established a CD8-targeted LNP delivery platform, we next sought to introduce cell-intrinsic control over CAR expression and overcome residual expression in non-T-cell populations (Supplementary Fig. 1G). To this end, we developed T-trex (T-cell– transient restricted expression), a modified mRNA strategy to restrict transgene expression to T cells and enhance functional potency of payloads.

To assess specificity, we first evaluated CAR expression across a panel of cell line models. Representative flow cytometry analyses demonstrated robust expression in PBMC-derived CD3⁺ T cells, with minimal to no detectable expression in non-T-cell lines (Fig. 2A). Transfection analysis of PBMCs further confirmed a high degree of T-cell restriction relative to non-T-cell populations (Fig. 2B), indicating that T-trex provides a layer of specificity that is independent of delivery bias alone.

**Figure 2.**
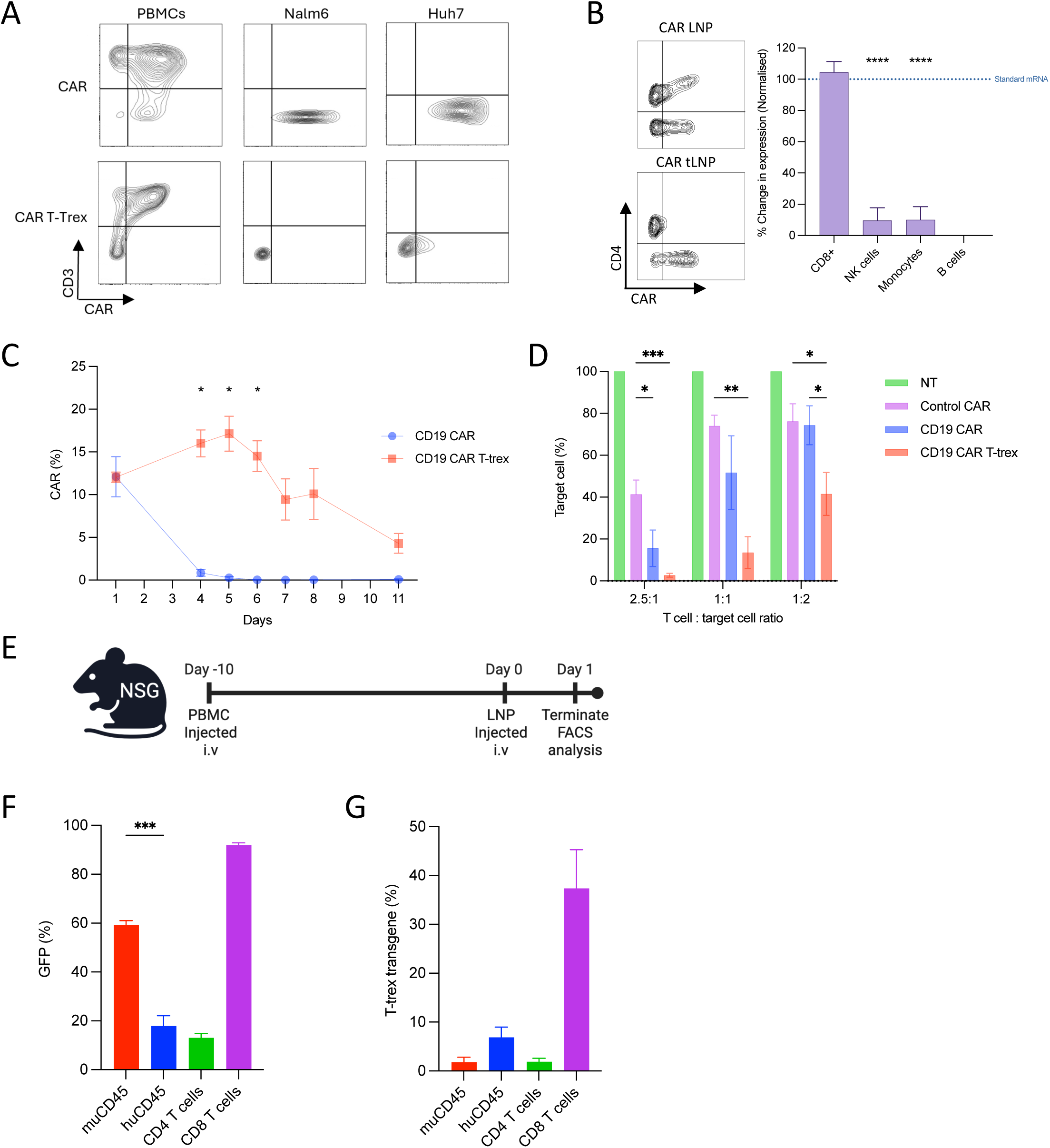
RNA engineering restricts expression to T cells and decouples CAR and IL-12 kinetics. A. Lineage-restricted expression of T-trex: Representative flow cytometry plots showing CAR mRNA expression in human PBMCs, Nalm6 (B-cell line), and Huh7 (liver-cell line). CAR mRNA exhibits non-specific expression across all cell lines, the CAR T-trex mRNA shows selective expression in CD3^+^ T cells. B. Quantitative analysis of expression specificity. Representative flow cytometry plots showing CAR mRNA expression in T cells, delivered using targeted and non-targeted LNPs (left panel). Graph showing normalised expression of T-trex mRNA in PBMC populations (left panel). tLNP delivers mRNA to CD8 cells, residual expression in non-T cells is minimised by using T-trex mRNA. n = 6 donors. Statistical significance was determined by two-way ANOVA. ****P<0.0001. C. Comparative expression kinetics. Longitudinal analysis of CAR expression following mRNA delivery in T cells. T-trex CAR mRNA (red) exhibits prolonged persistence compared to conventional CAR mRNA (blue). n = 3 donors. Statistical significance was determined by two-way ANOVA. *P<0.05. D. Antigen-specific cytotoxicity. *In vitro* cytotoxicity assay against CD19+ Nalm6 target cells at varying effector-to-target (E:T) ratios. CD19 CAR T-trex (orange) mediate potent killing that exceeds the efficacy of conventional CAR constructs (blue) and control benchmarks. n = 3 donors. Statistical significance was determined by two-way ANOVA. *P<0.05, **P<0.01, ***P<0.001. E. *In vivo* specificity study design. Timeline for evaluating RNA-level restriction in PBMC engrafted NSG mice. F. Quantification of unrestricted GFP mRNA *in vivo* following targeted LNP delivery. GFP expression was detected in engrafted human CD8 T cells and murine CD45 cells highlighting detectable off-target signals. n = 2 mice. Statistical significance was determined by one-way ANOVA. ***P<0.001. G. Quantification of T cell restricted, T-trex mRNA *in vivo* following targeted LNP delivery. T-trex mRNA construct demonstrate no detectable off-target expression in murine cells. n = 2 mice.

We next compared expression kinetics of a conventional CD19-CD28z CAR with its T-trex-encoded counterpart. CAR expression driven by the T-trex architecture was sustained over time and showed markedly prolonged persistence relative to the conventional CAR format (Fig. 2C). Mean fluorescence intensity analysis further demonstrated higher and more sustained surface expression of the T-trex-encoded CAR (Supplementary Fig. 2A), indicating that this approach enhances both the magnitude and duration of CAR expression. Although the precise mechanisms underlying this difference were not directly interrogated, the observed kinetics are consistent with slower turnover of T-trex CAR compared with conventional CARs, which may undergo tonic signalling–associated internalisation and dilution during T-cell proliferation.

To determine whether RNA-level restriction impacted functional activity, we assessed cytotoxicity against CD19⁺ Nalm6 target cells. T-trex-encoded CD19 CARs mediated potent antigen-specific killing that exceeded that observed with the conventional CAR construct and with previously reported *in vivo* CAR benchmarks^10^ (Fig. 2D). No cytotoxic activity was observed against CD19-negative SupT1 target cells (Supplementary Fig. 2B), confirming preservation of antigen specificity.

Finally, we assessed T-trex specificity *in vivo* (Fig. 2E). Following systemic administration of unrestricted GFP reporter mRNA, selective expression in CD8 T cells following CD8-targeted delivery was seen, but detectable off-target expression in murine CD45⁺ cells was also exhibited (Fig. 2F). Whereas delivery of a construct encoding the T-trex architecture showed no off-target signal, consistent with the combined effects of CD8-targeted delivery and RNA-level restriction (Fig 2G). Demonstrating that T-trex mRNA constructs embed an additional layer of restriction to combat residual expression in non-T cells.

Together, these data demonstrate that T-trex enables T-cell-restricted CAR expression while enhancing expression durability and functional potency. This architecture provides an additional control layer to targeted LNP delivery, supporting the development of dose-efficient and programmable *in vivo* CAR T cell therapies.

### Localised and transient IL-12 armouring enhances CAR potency while preserving antigen dependence

Having established delivery- and RNA-level control over CAR expression, we next sought to leverage this design to enhance the functional potency of *in vivo*–generated CAR T cells without increasing systemic exposure or compromising specificity. To this end, we engineered a T-cell-restricted (T-trex), surface-tethered interleukin-12 (IL-12) payload designed to provide local cytokine signalling in the context of CAR engagement, in contrast to conventional secreted cytokine delivery, which has been associated with dose-limiting toxicity ^13^ (Fig. 3A).

**Figure 3.**
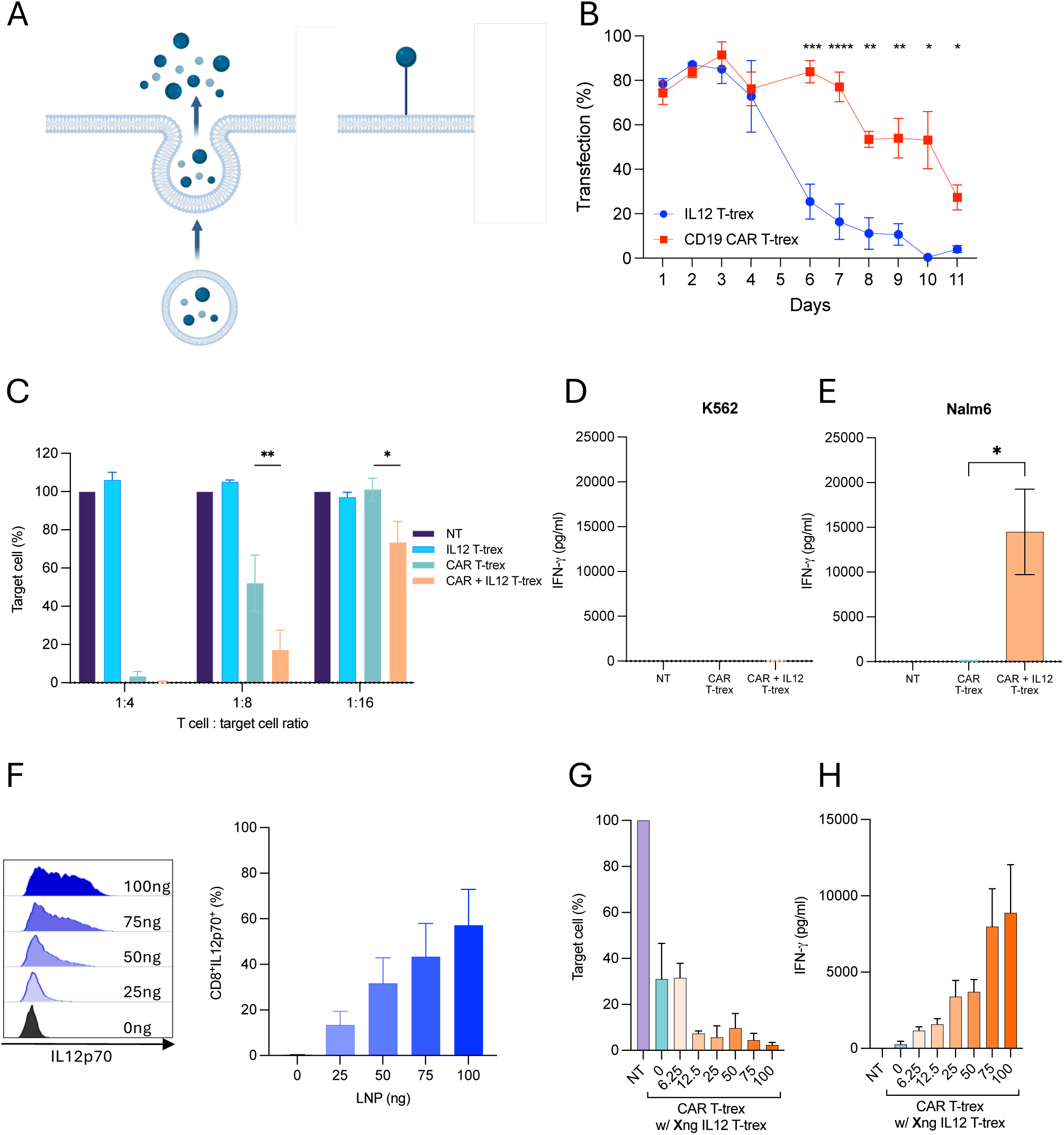
Surface-tethered IL-12 armouring lowers the functional activation threshold of CAR T cells. A. Schematic of surface-tethered IL-12 armouring. Comparison between conventional secreted cytokine delivery (left), which leads to systemic exposure and the surface tethered design (right). In this strategy, surface-tethered IL-12 provides localized signal amplification. B. Decoupled expression kinetics. Longitudinal analysis of surface CAR and IL-12 expression in primary human T cells following mRNA delivery. While both payloads utilise the T-trex architecture, the IL-12 armour (blue) exhibits a more transient profile compared to the CAR (red), consistent with a time-limited functional boost. n = 6 donors. Statistical significance was determined by two-way ANOVA. *P<0.05, **P<0.01, ***P<0.001, P<0.0001. C. Sustained cytotoxicity via IL-12 amplification. *In vitro* cytotoxicity against CD19^+^ Nalm6 targets at various effector-to-target (E:T) ratios measured at day 7 post-transfection. CAR armoured with tethered IL-12 (orange) maintains potent killing at lower E:T ratios where CAR alone (light blue) fails to sustain activity. n = 3 donors. Statistical significance was determined by a two-tailed Student’s t-test *P < 0.05, **P < 0.01. D–E. Antigen-dependent IFN-γ production. IFN-γ levels measured in co-cultures with antigen-negative K562 cells (D) or antigen-positive Nalm6 cells (E). IL-12 armouring dramatically increases IFN-γ production only in the presence of antigen, confirming that armouring preserves tight specificity and does not drive non-specific activation. n = 3 donors. Statistical significance was determined by a two-tailed Student’s t-test *P < 0.05. F. Tunable IL-12 expression. Representative flow cytometry histograms (left) and quantitative analysis (right) showing dose-dependent increases in surface IL-12p70 expression on CD8+ T cells following titration of IL-12-encoding LNPs. n = 3 donors. G–H. Titration of functional rescue. Impact of increasing IL-12 doses on CAR-mediated target cell killing (G) and corresponding IFN-γ secretion (H). Functional potency can be precisely titrated through LNP dose, allowing for a balanced optimization of cytotoxicity and cytokine output. n = 3 donors.

Expression kinetics of T-trex-encoded, tethered IL-12 were first evaluated following mRNA delivery. Representative flow cytometry analyses demonstrated detectable IL-12 expression shortly after transfection, followed by a decline over time (Supplementary Fig. 3A). Quantitative analysis confirmed that IL-12 expression was transient and time-limited, with a shorter duration of expression relative to the CAR payload (Fig. 3B), consistent with the intended armouring profile. *In vivo* assessment further supported restricted detection of IL-12 following systemic administration (Supplementary Fig. 3B), indicating that tethering combined with T-trex-mediated RNA-level control constrains cytokine exposure.

We next assessed the functional impact of IL-12 armouring on CAR-mediated cytotoxicity. In co-culture assays with CD19⁺ Nalm6 target cells, CAR T cells armoured with tethered IL-12 exhibited enhanced cytotoxic activity compared with CAR alone at later time points (day 7), indicating sustained functional benefit (Fig. 3C). Importantly, no cytotoxic activity was observed against antigen-negative target cells at early or late time points (Supplementary Figs. 3C and 3E), and enhanced killing was restricted to antigen-positive targets (Supplementary Fig. 3D), confirming preservation of antigen specificity.

To further characterise cytokine release profiles, IFN-γ production was measured during co-culture assays at a low effector-to-target ratio (1:8). In the presence of antigen-negative target cells, IFN-γ levels remained low for both CAR and CAR + IL-12 conditions (Fig. 3D), indicating that IL-12 armouring does not drive non-specific activation. In contrast, IFN-γ production was increased in the presence of antigen-positive target cells when IL-12 was co-expressed with the CAR (Fig. 3E), consistent with antigen-dependent amplification of CAR T-cell activity. Control conditions using non-transfected cells or IL-12 alone did not result in appreciable IFN-γ production in the presence or absence of antigen (Supplementary Figs. 3F and 3G), confirming that enhanced cytokine release requires CAR engagement.

We next examined whether IL-12 armouring could be titrated to balance potency and cytokine output. Transfection with increasing concentrations of IL-12-encoding LNP resulted in dose-dependent IL-12p70 expression, as assessed by flow cytometry (Fig. 3F). Correspondingly, CAR-mediated cytotoxicity against antigen-positive target cells increased with IL-12 dose (Fig. 3G), while IFN-γ production measured from the same co-cultures showed a proportional increase (Fig. 3H). These data demonstrate that IL-12 armouring is tunable and can be adjusted to enhance CAR potency while maintaining control over cytokine production.

Together, these results show that T-cell-restricted, tethered IL-12 functions as a local and transient amplifier of CAR T-cell activity. By enhancing antigen-dependent cytotoxicity without inducing non-specific activation, this armouring strategy provides a means to lower the effective activation threshold of *in vivo*–generated CAR T cells while preserving specificity and controllability.

### IL-12 armouring enables dose-efficient and repeatable in vivo CAR activity

To determine whether targeted delivery, RNA-level restriction, and IL-12 armouring could translate into meaningful improvements in *in vivo* efficacy, we next evaluated CAR activity across a series of dose-ranging and repeat-dosing studies (Fig. 4A). These experiments were designed to explicitly identify the dose limitations of unarmoured *in vivo* CAR delivery and to assess whether IL-12 armouring could overcome these constraints without compromising tolerability.

**Figure 4.**
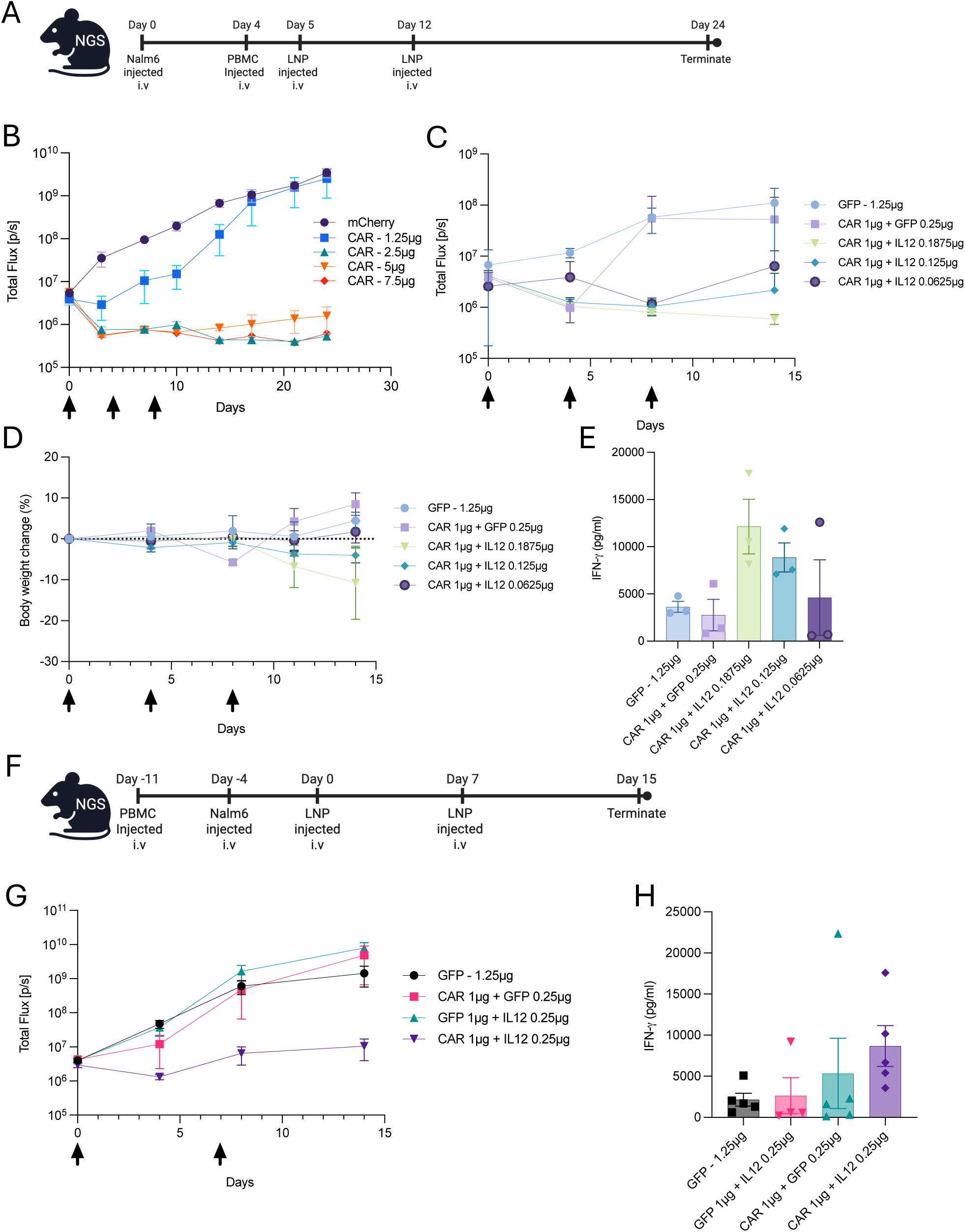
Repeat-dose *in vivo* CAR-T achieves sustained B-cell depletion at ultra-low total dose. A. Experimental study design B. Identification of suboptimal CAR dose. Longitudinal titration of unarmoured CD19 CAR mRNA (Total Flux [p/s]) identifies 1.25 µg as the suboptimal dose threshold at which CAR activity fails to sustain long-term tumour control. n = 3 mice. C. Functional rescue via tethered IL-12 titration. Evaluation of a fixed suboptimal CAR dose (1 µg) combined with increasing amounts of tethered IL-12 mRNA (titrated from 0.0625 to 0.1875 µg). Co-administration of IL-12 restores tumour control in a dose-dependent manner. n = 3 mice. D. Tolerability of dose titrations. Longitudinal body weight change (%) monitoring confirms that the various CAR and IL-12 dose combinations were well tolerated without significant adverse mass loss. n = 3 mice. E. Dose dependent cytokine production. Quantification of serum IFN-g in mice demonstrates a positive correlation between IL-12 mRNA dose and cytokine production. n = 3 mice. F. Repeat-dosing regimen schedule. Experimental timeline for evaluating LNP administration at extended seven-day intervals (Day 0 and Day 7) to assess the feasibility of maintaining disease control at low total doses. G. Sustained efficacy under repeat dosing. BLI quantification demonstrating that the armoured CAR (1 µg CAR + 0.25 µg IL-12) maintains robust and sustained tumour depletion compared to GFP or unarmoured controls when administered on an extended schedule. n = 5 mice. H. Serum cytokine. Serum IFN-g measured from mice demonstrates elevated cytokine production when CAR + IL-12 are administered, correlating with observed sustained efficacy. n = 5 mice.

We first performed a CAR dose titration using bioluminescence imaging (BLI) to monitor B cell tumour burden *in vivo*. At higher CAR doses, robust tumour control was observed; however, titration identified a suboptimal dose of 1.25 µg at which CAR mRNA alone failed to sustain antitumour activity (Fig. 4B; Supplementary Fig. 4A). Body weight monitoring demonstrated that CAR administration was well tolerated across the tested dose range (Supplementary Fig. 4B).

We next assessed whether IL-12 armouring could restore efficacy at this suboptimal CAR dose. Co-administration of tethered IL-12 with CAR mRNA resulted in a marked improvement in tumour control at the previously ineffective 1.25 µg CAR dose (Supplementary Fig. 4C), without inducing adverse changes in body weight (Supplementary Fig. 4D). These findings demonstrate that IL-12 armouring can functionally rescue CAR activity under dose-limited conditions.

To further explore the relationship between IL-12 dose and *in vivo* efficacy, we performed an IL-12 titration study in which a fixed CAR dose was combined with increasing amounts of IL-12-encoding mRNA. BLI analysis revealed a dose-dependent enhancement of tumour control with increasing IL-12 levels (Fig. 4C), while body weight measurements remained stable across the tested range (Fig. 4D). Serum cytokine analysis from this study revealed an IL-12 dose dependent IFN-g response (Fig. 4E), supporting the tuneability of IL-12 to modulate CAR function and cytokine responses.

Having established that IL-12 armouring enables robust efficacy at reduced CAR doses, we next asked whether this increased potency could support less frequent dosing. A repeat-dosing regimen was therefore evaluated performing LNP administration at seven-day intervals (Fig. 4F). Under this extended dosing schedule, sustained tumour control was maintained despite the low total CAR dose delivered (Fig. 4G). Analysis of serum cytokines showed elevated levels of IFN-g when CAR T-trex and IL-12 T-trex mRNAs are co-administered compared to CAR alone (Fig. 4H). Importantly, body weight remained stable throughout the repeat-dosing period (Supplementary Fig. 4E). Collectively, this demonstrates the ability of IL-12 to enhance CAR activity without causing toxicity.

## Discussion

The emergence of *in vivo* CAR T cell generation offers a path to overcome the manufacturing complexity, cost, and scalability limitations of *ex vivo*–engineered cell therapies. Recent studies have shown that systemically delivered CAR-encoding mRNA can reprogram T cells *in vivo*, resulting in antitumour activity in preclinical models and early clinical trials ^10,11^. However, current *in vivo* CAR approaches remain constrained by a fundamental dose–durability trade-off: CAR expression is transient, while dose escalation or frequent administration is restricted by toxicity and clinical feasibility.

Here, we describe a programmable *in vivo* CAR platform that addresses these limitations by enhancing the potency and duration of response. We have developed a novel LNP *in vivo* CAR platform which integrates targeted delivery, RNA-level expression gating, and local cytokine armouring to enhance the functional potency of CAR-expressing T cells. This integrated approach enables a safer and robust *in vivo* activity at substantially reduced total CAR exposure and supports repeat dosing at extended intervals.

A central component of this platform is the T-trex design, which restricts CAR expression to T cells. While prior *in vivo* CAR studies have relied primarily on delivery bias to favour T cell transfection, residual expression in non-T-cell compartments has remained a concern ^9,10^. By introducing RNA-level gating that operates independently of delivery chemistry, T-trex provides an additional and durable layer of specificity. Notably, T-trex–encoded CARs exhibited enhanced expression durability and functional potency. The observed kinetics are consistent with slower turnover compared with conventional CAR architectures, which are susceptible to tonic signalling, internalisation, and dilution during T cell proliferation ^14,15^.

Building on this foundation, we incorporated a transient, surface-tethered T-trex interleukin-12 (IL-12) payload to further lower the activation threshold of *in vivo*–generated CAR T cells. IL-12 is a potent immunomodulatory cytokine, but its clinical utility has been limited by systemic toxicity when delivered in secreted or constitutive formats ^13,16^. By restricting IL-12 expression to T cells, tethering it to the T-cell surface, and constraining its expression temporally, the platform enables local cytokine signalling in the context of antigen engagement while minimising systemic exposure. Functionally, IL-12 armouring enhanced antigen-dependent cytotoxicity and IFN-γ production without inducing non-specific activation, and these effects were tunable across a defined dose range ^17^.

Importantly, these design elements translated into a distinct efficacy profile *in vivo*. Dose-ranging studies demonstrated that unarmoured CAR mRNA delivery fails to sustain activity below a defined threshold, reflecting insufficient biological potency rather than dose-limiting toxicity. In contrast, IL-12 armouring restored efficacy at sub-threshold CAR doses and enabled sustained tumour control under repeat-dosing regimens with extended intervals. This was achieved using total CAR doses substantially lower than those reported in prior *in vivo* CAR studies ^10,11^, supporting a shift from dose-intensive to potency-driven therapy.

Although this study focused on CD19 and IL-12 as a model antigen, the platform principles described here are not antigen or cytokine armour-specific and may be broadly applicable to malignancies characterised by high tumour burden, antigen re-emergence, or the need for sustained immune pressure. In such settings, the ability to administer CAR therapies repeatedly at low doses may enable dynamic control of effector function while limiting toxicity. Beyond oncology, similar principles may also apply to immune-mediated diseases requiring durable target cell depletion without prolonged CAR persistence.

Several limitations warrant consideration. A critical consideration for any repeat-dosing strategy is the potential for neutralizing anti-drug antibodies (ADAs) against the targeting ligand. To mitigate this, we utilized the AbNatiV deep learning platform to engineer VHH binders indistinguishable from human immune-system-derived sequences. Unlike standard humanisation which can destabilize the VHH fold, this approach optimises specific ’nativeness’ scores to maintain solubility and stability, key parameters for efficient targeted LNP formulation^12,18^.

In summary, this work establishes a programmable *in vivo* CAR T cell platform that integrates targeted delivery, RNA-level expression gating, and transient cytokine armouring to achieve dose-efficient and repeatable therapeutic activity. By shifting the design paradigm from delivery efficiency alone to functional potency and control, our platform provides a framework for next-generation *in vivo* CAR therapies capable of addressing key limitations that have constrained the field to date.

## Methods

### Anti-CD8 VHH binder production and characterisation

Ten anti-human CD8 VHH binders were developed by optimising previously described anti-CD8 VH and VL sequences with rational design strategies. Designed binder sequences were then C-terminally tagged with Cys-8xHis, synthesized and constructed into pTT5 expression vector (termed as CMRP784 – 794).

The constructed binders were then expressed in HEK293 cells for 7 days, and expressed binders were purified from the cleared conditioned medium with a Ni-Excel column (Cytiva). Purified binders were then buffer changed into PBS by dialysis, reduced with TCEP (5:1 molar ratio), and then dimerized with DHAA (50:1 molar ratio). Dimerized binders were further purified with a Ni-Excel column followed by a Superdex 200 size exclusion column (Cytiva). The purity of binders was determined with SDS-PAGE under reduced or non-reduced conditions. Final purified dimer binders were concentrated with a TFF cassette, aliquoted and stored at – 80 °C.

The affinity of the binders to human CD8 protein was analysed on a Biacore T200 instrument (Cytiva) using a Protein A sensor chip, following the manufacturer’s instructions. In brief, Fc-tagged human CD8 protein was captured at 5 µg/mL onto Flow Cell 2 (FC2) for 30 s at a flow rate of 10 µL/min in HBS-EP buffer (10 mM HEPES, 150 mM NaCl, 3 mM EDTA, 0.005% Tween-20, pH 7.4). Flow Cell 1 (FC1) served as a reference. After a 30 s stabilization phase, analytes were injected at concentrations starting at 100 nM with a 1:2 serial dilution series. Association was monitored for 120 s and dissociation for up to 1600 s at a flow rate of 30 µL/min over both flow cells. A double blank (buffer only) was included for reference subtraction, and one analyte concentration was run in duplicate as a control for reproducibility. Surfaces were regenerated using 30 s injections of glycine-HCl (pH 1.5) at 30 µL/min, repeated twice per cycle.

Thermal stability of the binders was analyzed with a Tycho NT.6 instrument (NanoTemper Technologies), following the manufacturer’s instructions. In brief, binders diluted to 1 mg/ml were loaded onto the instrument and intrinsic fluorescence at 330 nm and 350 nm was measured during a temperature ramp from 35 °C to 95 °C at 30 °C/min. The inflection temperature (T_i_), indicating thermal unfolding, was calculated automatically from the 350/330 nm fluorescence ratio. T_i_ values were used to compare relative stability between proteins, with measurements performed in duplicate to ensure reproducibility.

Functional testing of the binders was measured by their ability to enhance the transfection primary human CD8 T cells when being conjugated to the surface of LNP (LNP production and VHH conjugation processes are detailed in the sections below). LNPs encapsulated with eGFP mRNA (GenScript) were either unconjugated (control) or conjugated with different anti-human CD8 VHH binders were serially diluted and incubated with 1.5 × 10⁵ human T cells in RPMI1640 (Thermo Fisher Scientific) supplemented with 10% FBS (Labtech, Heathfield, UK) in 96-well U bottom well plates and cultured at 37 °C for 24 hours. After incubation, cells were washed and stained with APC-hCD4 (BioLegend) and DAPI (Thermo Fisher Scientific), and then eGFP expression on CD4+ and CD4-populations were evaluated on a MACSQuant® 10 flow cytometer (Miltenyi Biotec).

### mRNA synthesis and preparation

mRNAs encoding for eGFP, firefly luciferase and mCherry were purchased from GenScript, ProBio and CATUG, respectively. All other mRNAs used in this study were designed in-house, constructed into a modified pcDNA3.1 vector (Thermo Fisher Scientific) carrying a 5’ UTR, 3’ UTR and a 40A(G)40A polyA tail that can be transcribed from an upstream T7 promoter.

Small scale mRNA production was made in-house for in vitro studies. These mRNAs were made using a HiScribe® T7 mRNA Kit with CleanCap® Reagent AG (New England BioLabs), according to the manufacturer’s instructions. In brief, plasmids were first linearized with BspQI (New England BioLabs) and cleaned with a Monarch® Spin PCR & DNA Cleanup Kit (New England BioLabs), and in vitro transcribed into mRNA using HiScribe® T7 mRNA Kit with CleanCap® Reagent AG with 100% substitution of UTP with N1-Methyl-Pseudo-UTP. After in vitro transcription (IVT), template was moved by DNaseI treatment and mRNA was purified by LiCl precipitation and 70% ethanol wash. Final mRNA was dissolved in RNA storage buffer (Thermo Fisher Scientific) and stored at – 80 °C.

Large scale mRNA productions for *in vivo* studies were made by either GenScript or ProBio. These mRNAs were made with the manufacturers’ proprietary IVT protocols, purified with ion exchange columns, filtered to remove bacteria, concentrated with TFF cassettes. These mRNAs all have the same 100% substitution of UTP with N1-Methyl-Pseudo-UTP and a Cap1(3’-OMe)-AG cap.

### LNP production and VHH conjugation

A proprietary LNP formulation was obtained from an external supplier and used in the current study. mRNAs were encapsulated into LNPs using either NanoAssemblr Ignite or Blaze nanoparticle formulation system (Cytiva). Formulated materials were then diluted 10 folds into PBS (Thermo Fisher Scientific), concentrated and buffer-exchanged into LNP storage buffer with either 100 kD cut-off Amicon ultrafiltration columns (Merck) or 30 kD cut-off TFF cassettes (Cytiva). RNA encapsulation efficiency was measured with Quant-iT

RiboGreen Kit (Thermo Fisher Scientific) according to the manufacturer’s protocol. Particle size and polydispersity (PDI) were measured by Zetasizer Ultra (Malvern Panalytical), according to the manufacturer’s instructions.

For VHH conjugation onto the LNP surface through the maleimide-thiol chemistry, purified anti-human CD8 VHH dimers were first reduced with TCEP and then mixed with LNP at various ratios at room temperature for 1-2 h. Conjugated LNPs (tLNPs) were then treated with L-cysteine to quench extra reactive maleimide and conjugates were purified using 20 nm cut-off qEV size exclusion columns (IZON), following the manufacturer’s instructions.

Conjugation of VHH on the LNP surface was confirmed by NanoFCM NanoAnalyzer. tLNPs were first incubated with Alexa Fluor™ 488-human CD8 protein for 2 h at room temperature, and then samples were diluted 100 folds and evaluated on NanoFCM NanoAnalyzer, following the manufacturer’s instructions.

### Primary cells and cell culture

Peripheral Blood Mononuclear Cells (PBMCs) were isolated from whole blood by density centrifugation via Ficoll-Paque PLUS (GE-Healthcare) according to manufacturer’s protocol. Isolated PBMCs were cultured in complete media, constituting of RPMI 1640 (GIBCO) supplemented with 10% fetal bovine serum (FBS) (Labtech) and 2mM GlutaMAX (Gibco)

Nalm6 (DMSZ), K562 (ATCC) and SupT1 (ATCC) cells were cultured in complete media. HUH-7 (Sigma Aldrich) cells were cultures in Dulbecco’s modified eagle medium (GIBCO) supplemented with 10% FBS and 2mM GlutaMAX.

### Transfection

Primary cells and cell lines were harvested and resuspended in complete media with 10 ng/mL IL-2 (Miltenyi Biotec) at a density of 0.5-1×10^6^ cells/mL. LNP formulations were added to the cell suspension at a range of concentrations from 0.625-100ng. Following a 4-hour incubation at 37 °C and 5% CO_2_, cells were washed twice with sterile PBS (GIBCO) to remove unbound LNPs. The transfected cells were then transferred into complete media supplemented with 10 ng/mL IL-2 for experimentation.

### Flow cytometry

Cells were stained with fluorescence-labelled antibodies for 20 minutes in subdued light with antibodies diluted in cell staining buffer (420201, Biolegend). Cells were stained with anti-mCD45-BV605 (clone 30-F11, #563053, BD Biosciences), anti-hCD45-PE-Cy7 (clone 2D1, #570764, BD Biosciences), anti-hCD19CAR detection reagent-Alexa Fluor 647 (clone Y4S, #FM3-AM534-200, Acro Biosystems), anti-hCD3-Per-Cp-Cy5.5 (clone SK7, #344808, Biolegend), anti-hCD4-APC-Cy7 (clone RPA-T4, #557871, BD Biosciences), anti-hCD20-PE Texas Red (clone HI47, #MHCD2017, ThermoFisher Scientific), anti-G4S linker – PE (clone 016, #G4S-PFMY25, Acro Biosystems), IL12p70-PE (clone 20C2, #559325, BD Biosciences), DAPI (4’,6-Diamidino-2-Phenylindole, Dilactate) – (#422801, Biolegend) and CountBright Absolute Counting beads (Life Technologies; C36950). Flow cytometry was performed using the LSR Fortessa (BD Biosciences) and analysed using FlowJo version 10 software (BD Biosciences).

### Time course studies

Transfected T cells are cultured in complete media in the complete media supplemented with 10ng/ml IL2 and maintained in culture to up to 11 days. Changes in expression are monitored at regular intervals. Between T-trex CAR and conventional CAR, expression was measured at days 1, 4, 5, 6, 7, 8 and 11 post-transfection. T-trex CAR and T-trex IL12 expression kinetics was assessed at days 1, 2, 3, 4, 6, 7, 8, 9, 10 and 11 post-transfection.

### Cytotoxicity assay

T cells are co-cultured with K562, Nalm6 or SupT1 target cells at various T cell : target ratios from 1:1, 1:4, 1:8 and 1:16, with target cells remaining constant at 5×10^4^. Target cells are enumerated via flow cytometry using counting beads and cytotoxicity calculated as a percentage by normalising to the number of target cells recovered from co-cultures with non-transfected (NT) T cells. Culture supernatants were harvested and assessed for presence of IFN-g via the ELISA MAX Deluxe set human IFN-g kit (Biolegend) according to the manufacturers protocol.

### In vivo studies

*In vivo* studies were carried out according to regulatory guidelines and conducted by Frontage Laboratories Inc (China) and Crown Bioscience (UK).

Biodistribution and expression studies were conducted using CD-1 mice and C57BL/6 mice (Charles River Laboratories). PBMC models were conducted using M-NSG (NOD.Cg-Prkdc^scid^Il2rg^em1Smoc^) mice (Shanghai Model Organism, China). All animals were raised under pathogen-free conditions.

### Luciferase biodistribution models

For biodistribution study in mouse, female C57BL/6J mice (Crown Biosciences) at 5 – 8 weeks old were first treated with PBS or LNP1 encapsulated with firefly luciferase (LNP1-fLuc) at a dose of 2ug through i.v. Four hours later, mice were injected (s.c.) with 150 mg/kg D-Luciferin and 10 minutes following administration, mice were anaesthetised for full body live imaging (IVIS Spectrum BL), and then animals were scarified and organs were harvested for a second round of imaging.

For biodistribution study in NHP, cynomolgus monkeys (Macaca fascicularis) (Hainan Jingang Co Ltd., China) of both sexes, weighing 3–8 kg, were pre-screened and confirmed negative for anti-PEG antibodies prior to study initiation. Animals were treated with Saline or LNP1-fLuc at a dose of 0.1 mg/kg through i.v. infusion. Six hours after treatment, animals were euthanized and major organs harvested and luciferase signal was quantified.

### PBMC model

Previously cryopreserved donor derived human PBMCs were thawed and 5×10^6^ cells/ animal was injected intravenously (i.v) via the tail vein into M-NSG (NOD.Cg-Prkdc^scid^Il2rg^em1Smoc^, Shanghai model organism, China) mice. After 7 days, blood was collected for analysis by flow cytometry and 1×10^6^ Nalm6 cells were injected i.v. Five days after inoculation, tumour presence was confirmed by bioluminescence imaging. The mice were randomized based on PBMC engraftment and tumour burden on day 0 and injected i.v with tLNPs. Mice were treated every 4 or 7 days receiving up to 3 doses. Changes in tumour burden and bodyweight was measured twice weekly over the duration of the study.

### Statistics

Data analysis was performed using GraphPad Prism 10 (Dotmatics, USA). Lines and error bars represent mean and standard error. Analysis was done using One-way ANOVA or Two-way ANOVA followed by two-tail Student t-test. Non-significance when P>0.05 is not highlighted. When significance is seen, annotations were denoted as: *P < 0.05; **P < 0.01; ***P < 0.001; ****P < 0.0001.

## Author contributions

S.O. and S.P.C. conceived and devised the study. S.O., S.S and K.H. wrote the manuscript. S.S. also contributed to study design.

K.H. and B.M. devised the targeting strategy and carried out lipid nanoparticle (LNP) formulation and testing.

Y.Y. performed in vitro and *in vivo* analysis of LNP formulations.

H.W. conducted in vitro analysis of LNP formulations.

J.H. performed binder characterisation studies.

M.M. and E.S. carried out in vitro characterisation of CAR constructs.

P.S., A.R., and M.G. developed computational models for binder design and performed humanisation and optimisation.

## Acknowledgements

This research was funded in part by Innovate UK (UK Research and Innovation -Grant number 10104367).

