## Supplementary figures for "Engineering armoured *in vivo* CAR T cells through targeted delivery and transient mRNA gating"

**Supplementary Figure 1.**

A. AbNatiV-based computational humanisation: Structural modelling and residue-level humanness profiling identify low-humanness non-CDR surface residues. Iterative sampling humanises these sites while monitoring CDR displacement, reducing predicted immunogenicity without loss of structural or physicochemical stability.

B. Biophysical characterisation of humanised CD8 binders: Binding affinities ($K_D$) and kinetics (ka, kd) were measured by surface plasmon resonance SPR against recombinant human CD8 protein. Stability was determined by assessing thermal inflection and melting ($T_i/T_m$) using the Tycho NT.6. Representative SPR sensorgrams show affinity and dissociation diversity for all candidates (CMRP784–794) across a 6.25–100 nM range.

C. Modular screening platform for rapid VHH evaluation: Schematic of the "molecular glue" strategy used for universal VHH capture. Unconjugated LNPs are converted into universal capture LNPs through incorporation of a molecular glue component. Targeted LNPs are subsequently generated by the covalent, irreversible capture of tagged VHH binders, enabling high-throughput evaluation of multiple CD8-targeting candidates.

D. Specificity of humanised VHH candidates in $CD4^+$ T cells. Representative: GFP transfection in $CD4^+$ T cells following treatment with $CD8^+$ VHH-targeted LNPs. All candidates exhibited negligible GFP expression in $CD4^+$ T cells across the concentration range, confirming lineage specificity for the selected binders.

E. TCEP treatment of the VHH dimers. Dimeric VHH was treated with TCEP at various concentrations for 1hr at room temperature, with a non-reducing SDS-PAGE gel to evaluate reduction efficiency.

F. Physical characterisation and surface decoration of targeted LNPs: (Top) Summary table of encapsulation efficiency, hydrodynamic size, polydispersity index (PDI), and VHH conjugation efficiency for untargeted (LNP1) and targeted (tLNP1) formulations. (Bottom left) Representative dynamic light scattering (DLS) size distribution profiles demonstrating a modest increase in hydrodynamic diameter upon VHH conjugation while maintaining a low PDI. (Bottom right) Nano Flow Cytometry-based analysis of LNP surface decoration, showing a shift in FITC signal for tLNP1 compared to LNP1, confirming ligand conjugation.

G. Lineage specificity across PBMC compartments. Quantitative analysis of GFP in cell subsets ($CD4^+$ T, $CD8^+$ T, B cells, monocytes and NK cells) from PBMCs following treatment with CD8-targeted tLNP1. n = 2 donors. Statistical significance was determined by one-way ANOVA. $**P<0.01$, $***P<0.001$.

**Supplementary Figure 2.**

A. Quantitative mean fluorescence intensity (MFI) analysis. Longitudinal MFI analysis demonstrating higher and more sustained surface expression levels of T-trex-encoded CARs (orange) relative to the conventional CAR format (blue) in primary human T cells. n = 3 donors. Statistical significance was determined by two-way ANOVA. **$P<0.01$, ***$P<0.001$.

B. Preservation of antigen specificity. *In vitro* cytotoxicity assay against CD19-negative SupT1 target cells at varying effector-to-target ratios. Neither conventional nor T-trex constructs mediated killing of antigen-negative cells, confirming that enhanced potency does not compromise specificity. n = 3 donors.

**Supplementary Figure 3.**

A. Longitudinal expression of IL-12 and CAR payloads. Representative flow cytometry plots showing the surface expression kinetics of T-trex-encoded, tethered IL-12 (top) and CD19 CAR (bottom) in primary human T cells over 10 days post-mRNA delivery. Expression of IL-12 is detectable shortly after transfection and follows a transient, time-limited profile, declining more rapidly than the associated CAR payload.

B. *In vivo* restriction of IL-12 expression. Representative flow cytometry plots analysing human $CD45^{+}$, human $CD45^{+}CD3^{+}$, and murine $CD45^{+}$ populations from PBMC engrafted NSG mice following systemic administration of CD8-targeted LNPs. The data demonstrate that IL-12 expression is strictly confined to the human T-cell compartment, with no detectable signal in murine cells, confirming that the combination of cell surface tethering and T-trex-mediated RNA-level control minimizes systemic exposure.

C–E. Specificity of cytotoxic activity. Percentage of target cells remaining in co-culture assays at various effector-to-target (T cell : target cell) ratios. (C, E) Killing assays against antigen-negative target cells at early (C) and late (E) time points post-transfection demonstrate no non-specific cytotoxicity from the armoured platform. (D) Killing assays against antigen-positive $CD19^{+}$ Nalm6 target cells, showing that CAR armoured with IL-12 significantly enhances cytotoxic activity compared to CAR alone or non-transfected controls. n = 3 donors. Statistical significance was determined by a two-tailed Student's t-test *$P < 0.05$.

F–G. Analysis of baseline cytokine production. IFN-γ secretion (pg/mL) measured from non-transfected (NT) or IL-12-only transfected T cells in the presence of antigen-negative K562 cells (F) or antigen-positive Nalm6 cells (G). Negligible cytokine production is observed for these controls in the absence of CAR-antigen engagement, confirming that enhanced cytokine release is antigen-dependent and requires CAR activation. n = 3 donors.

**Supplementary Figure 4.**

A. Individual tumour kinetics during CAR dose titration: Representative longitudinal bioluminescence (BLI) images of individual NSG mice engrafted with $CD19^{+}$ Nalm6 cells. Images track tumour burden over 24 days following systemic administration of untargeted mCherry (control) or increasing doses of unarmoured CD19 CAR mRNA (1.25 µg to 7.5 µg).

B. Tolerability of unarmoured CAR dose ranging: Longitudinal body weight change (%) monitoring for mice treated with the unarmoured CAR titration series. High-dose administration (7.5 µg) resulted in transient weight loss, while lower doses remained well-tolerated. n = 3 mice.

C. Functional rescue of suboptimal CAR activity: Quantitative BLI analysis (Total Flux [p/s]) comparing tumour burden in mice treated with a suboptimal dose of unarmoured CAR (1.25 µg) versus the same dose armoured with tethered IL-12 (1 µg CAR + 0.25 µg IL-12). Co-administration of IL-12 significantly improved tumour control at the previously ineffective CAR dose. n = 3 mice.

D. Tolerability of IL-12-mediated functional rescue: Longitudinal body weight monitoring for the groups in Panel C, confirming that functional rescue via surface-tethered IL-12 armouring is well-tolerated without inducing systemic cachexia. n = 3 mice.

E. Safety profile during repeat dosing: Body weight change (%) monitoring throughout the 7-day interval repeat-dosing regimen. Stable body mass across all treated groups indicates a lack of cumulative toxicity from repeated LNP administration. n = 5 mice.

Supplementary Figure 1

A

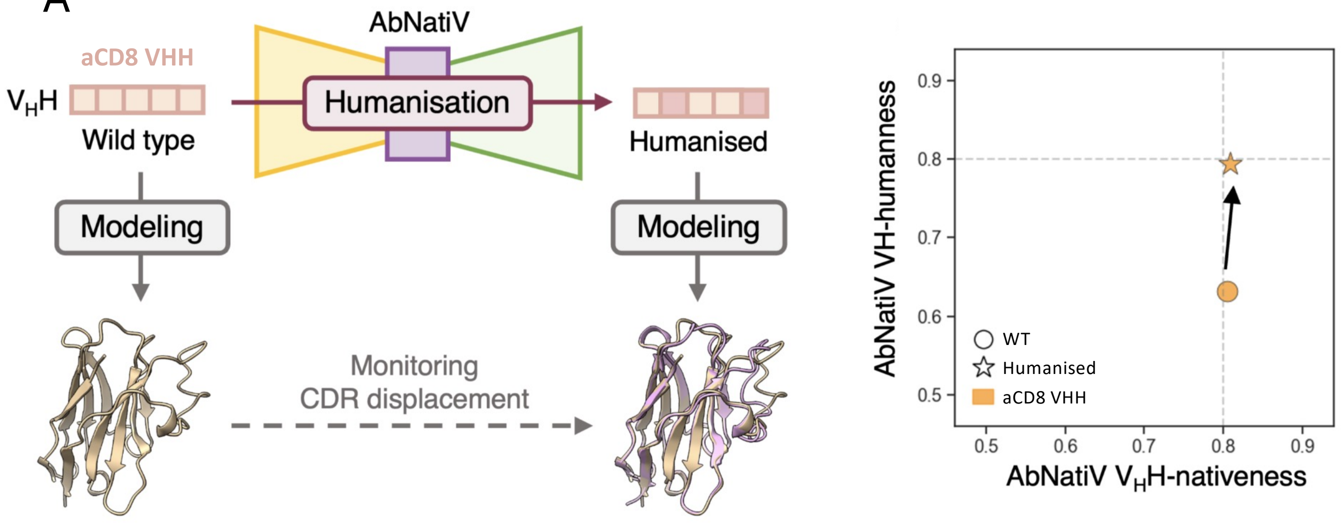


B

| Binder ID | Biacore KD against Human CD8 | | | | Thermostability (Tm) | |
|---|---|---|---|---|---|---|
| | ka (1/Ms) | kd (1/s) | KD (M) | KD (nM) | Mean | SD |
| CMRP784 | 3.83E+05 | 1.34E-04 | 3.49E-10 | 0.35 | 74.18 | 0.10 |
| CMRP785 | 3.77E+05 | 0.001591 | 4.22E-09 | 4.22 | 73.84 | 0.01 |
| CMRP786 | 3.22E+05 | 0.0029 | 9.01E-09 | 9.01 | 74.21 | 0.08 |
| CMRP787 | 8.81E+05 | 0.006423 | 7.29E-09 | 7.29 | 73.45 | 0.07 |
| CMRP788 | 2.68E+03 | 0.001987 | 1.26E-08 | 741.10 | 74.68 | 0.13 |
| CMRP789 | 1.55E+05 | 2.84E-04 | 1.83E-09 | 1.83 | 71.91 | 0.30 |
| CMRP790 | 3.01E+05 | 0.002555 | 8.49E-09 | 8.49 | 71.89 | 0.03 |
| CMRP791 | 2.01E+05 | 0.006167 | 3.07E-08 | 30.72 | 72.17 | 0.06 |
| CMRP792 | 1.57E+05 | 0.003972 | 2.54E-08 | 25.35 | 73.11 | 0.25 |
| CMRP794 | 2.17E+06 | 3.69E-04 | 1.70E-10 | 0.17 | 74.96 | 0.11 |

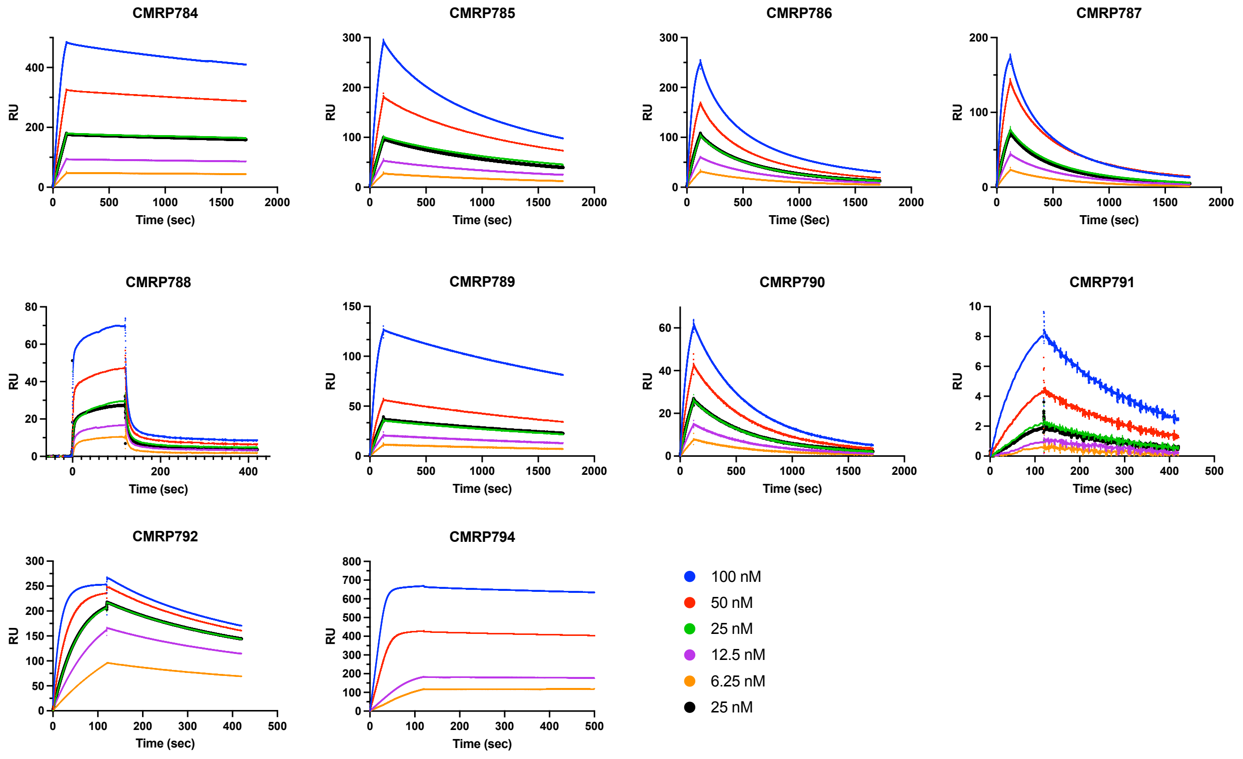

Supplementary Figure 1

C

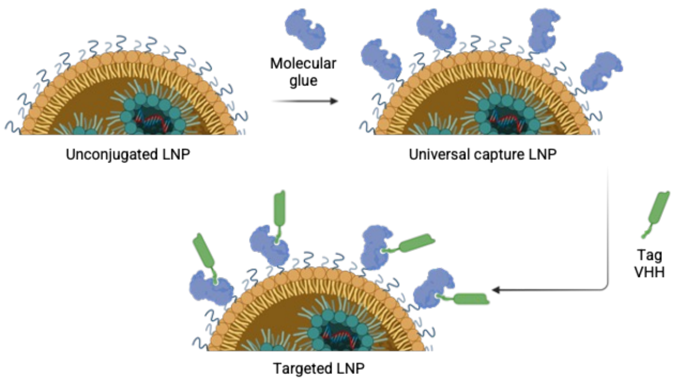


E

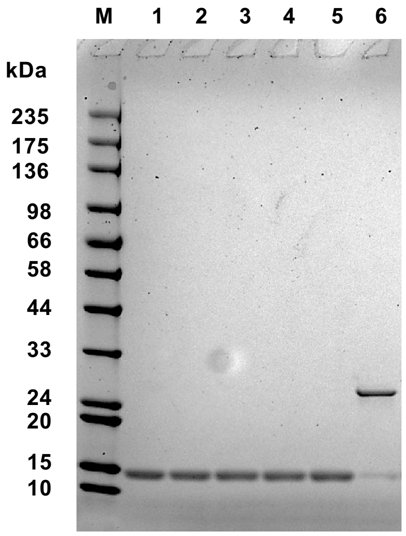


M: Marker
1. Dimeric VHH treated with 30 μM TCEP
2. Dimeric VHH treated with with 60 μM TCEP
3. Dimeric VHH treated with with 90 μM TCEP
4. Dimeric VHH treated with With 120 μM TCEP
5. Dimeric VHH treated with with 150 μM TCEP
6. Untreated dimeric VHH

D

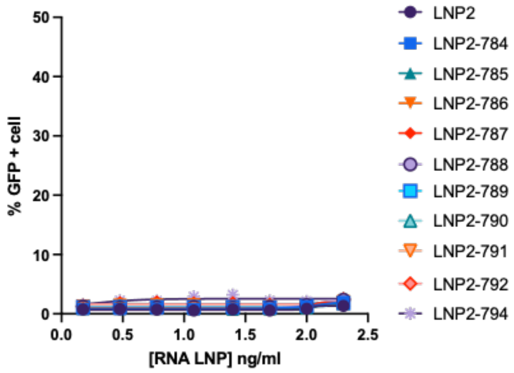


F

| Sample ID | EE (%) | Size (nm) | PDI | Conjugation (%) |
|---|---|---|---|---|
| LNP1 | 97.0 | 99.58 ± 1.29 | 0.072 ± 0.021 | n/a |
| tLNP1 | 96.2 | 108.3 ± 0.83 | 0.078 ± 0.012 | 82.6 ± 3.5 |

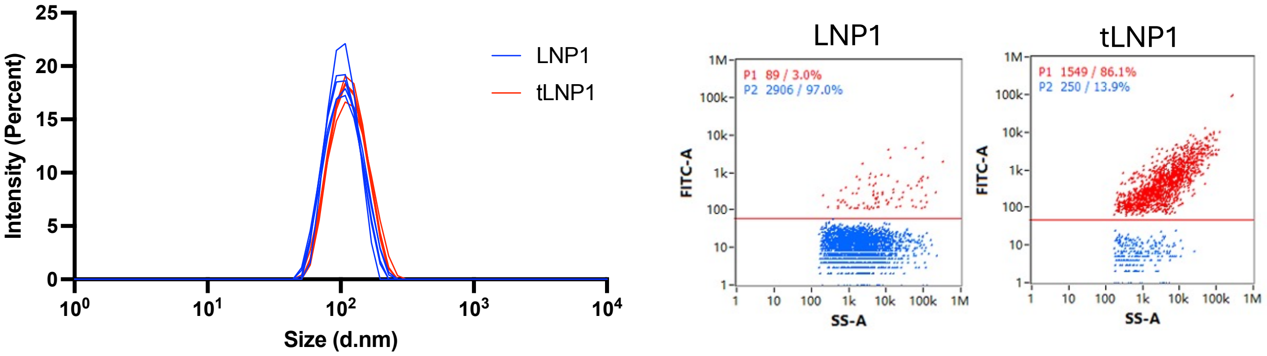


G

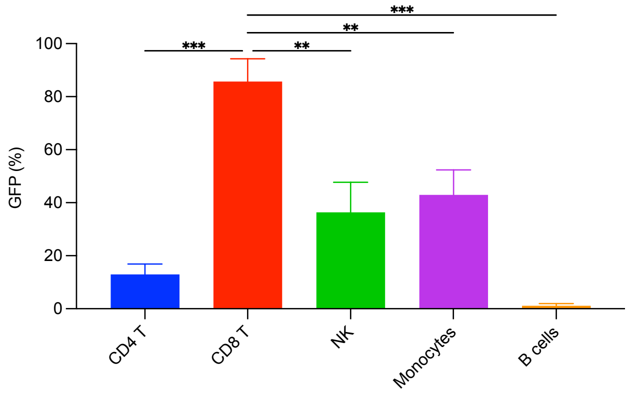

Supplementary Figure 2

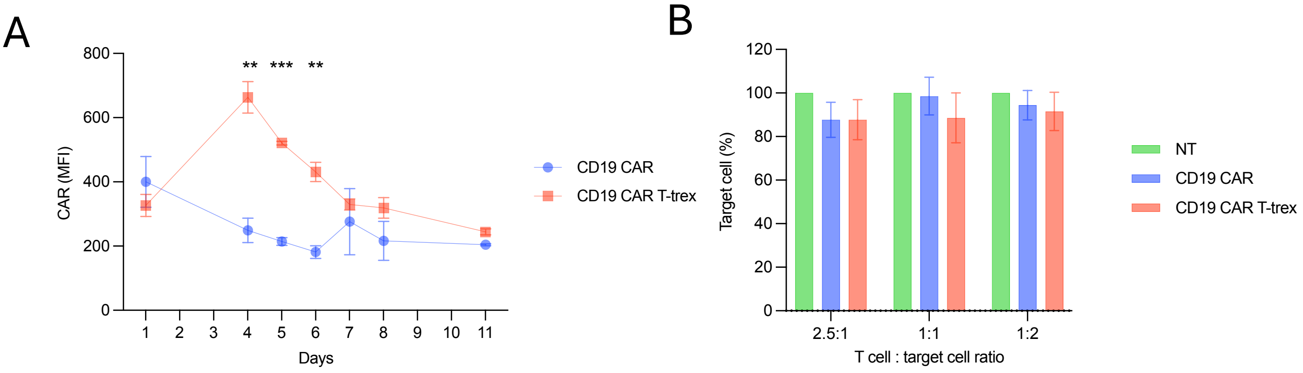

Supplementary Figure 3

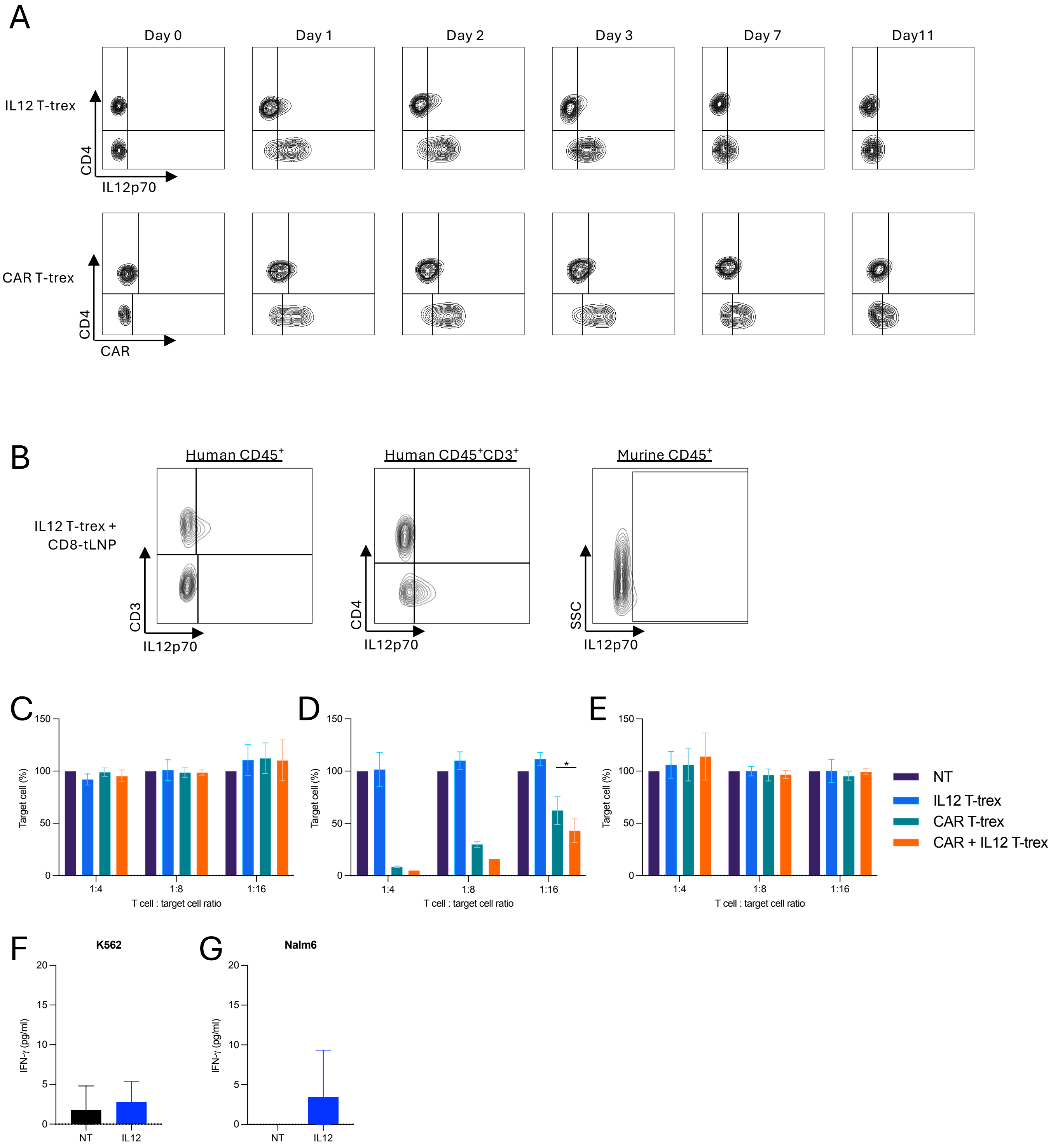

Supplementary Figure 4

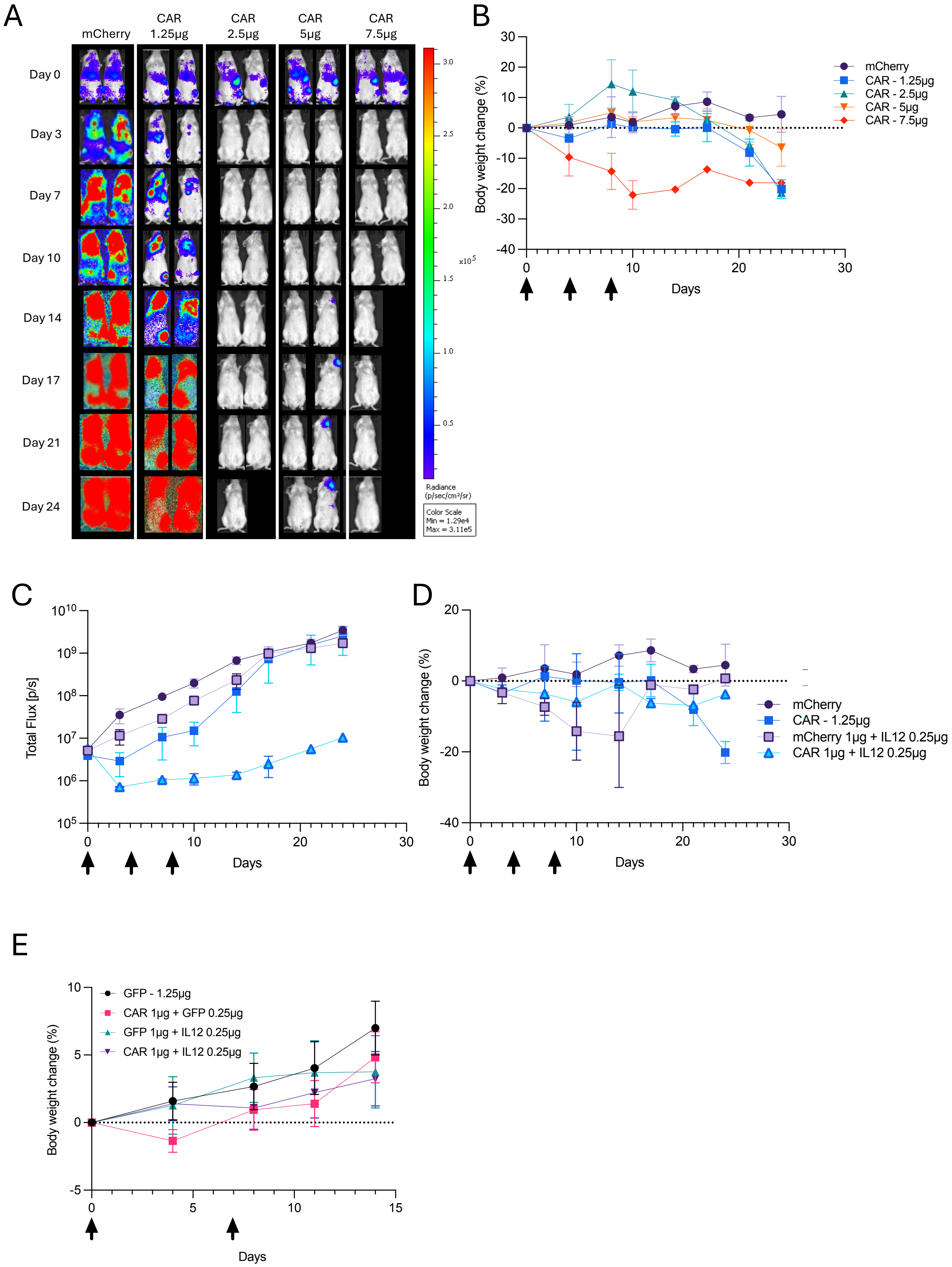
